# Cardiorespiratory fitness and sex impact the colorectal cancer-suppressive effects of aerobic exercise serum

**DOI:** 10.64898/2026.10.15.759428

**Authors:** Rachael N. Kemp, Ume-Kulsoom Shah, Sarah Alwan, David Oyler, Gareth Jenkins, Kelly A. Mackintosh, Melitta A. McNarry

**Author notes:** Corresponding author: Rachael Kemp.

## Abstract

Post-exercise human serum treatment can exhibit anti-viability properties in various cancer cell lines. However, whether the presence and/or magnitude of the effect differs in different participant populations is yet to be established. Here, we characterised baseline differences in serum-stimulated colorectal cancer cell viability and the acute exercise-induced response in males and females with varying levels of cardiorespiratory fitness (CRF). Forty-one participants (13 female) were stratified by maximal exercise test-derived V̇O_2_peak into high (*n* = 20) and low CRF (*n* = 21) and completed two experimental trials in a randomised crossover design: (1) 30-min moderate-intensity aerobic exercise; and (2) 30-min seated rest. Serum samples were collected before (pre), immediately after (post0), and 60-min after (post60) each trial and used to treat the human colorectal cancer cell line Caco-2 for 48 h, with cell viability assessed by MTT assay. Exercise-induced alterations in Caco-2 cell viability were, at least in part, modulated by CRF and sex. Specifically, at post0 and post60, exercise serum significantly reduced cell viability compared with no-exercise control serum when derived from individuals with high but not low CRF, and when derived from males but not females. These findings provide the first insights into the role of individual characteristics in determining the effects of exercise on colorectal cancer cells, with potential implications for increasing the precision of exercise recommendations for colorectal cancer risk reduction.

## Introduction

Colorectal cancer is the third most common cancer and second leading cause of cancer mortality globally (Bray e*t al*., 2024). While the aetiology of colorectal cancer is multifactorial, it is estimated that around half of cases are attributable to modifiable lifestyle risk factors (Brown *et al*., 2018). Accordingly, numerous epidemiological studies consistently demonstrate that higher levels of physical activity (McTiernan *et al*., 2019; Moore *et al*., 2016) and cardiorespiratory fitness (CRF; Bojsen-Møller *et al*., 2024; Hillreiner *et al*., 2020) are associated with a lower colorectal cancer risk. These associations may, at least in part, be attributed to differences in the circulating milieu, such as metabolic hormones, microRNAs, lower levels of pro-inflammatory cytokines, and enhanced immune surveillance (Kunutsor *et al*., 2024; Wang *et al*., 2022).

To investigate the direct effects of exposure to different circulating milieu on cancer biology, in vitro models can be used to culture cancer cell lines with ex vivo human serum or plasma (Cela *et al*., 2025). Whilst evidence is currently limited, cross-sectional studies conducted in a resting state suggest that serum/plasma derived from individuals with higher CRF reduces the viability of chronic lymphocytic leukaemia and prostate cancer cell lines compared with serum/plasma from those with lower CRF (Sitlinger *et al*., 2021; Sridhar *et al*., 2017), although others reported no differences (Peres *et al*., 2022). However, the impact of CRF on colorectal cancer cells remains unexplored. Additionally, no studies have investigated the interplay between CRF and non-modifiable risk factors, such as sex, on serum-stimulated cancer cell activity across any cancer type, despite the importance of such insights for developing more targeted cancer prevention strategies.

In addition to the basal circulating profile, there is growing interest in understanding the effects of acute physiological states, such as exercise, on cancer cells (Hojman *et al*., 2018; Dethlefsen e*t al*., 2017a). Treatment of various cancer cell lines with serum derived immediately after a single bout of exercise can reduce cell viability compared to pre-exercise or control serum (Metcalfe *et al*., 2021; Devin *et al*., 2019). However, a significant degree of variability in the effects of acute exercise-conditioned serum has been reported to date (Orange *et al*., 2020), which may be partly attributable to differences in participant populations. For instance, sex and training status have been suggested to influence the exercise-induced temporal pattern and magnitude of responses in numerous myokines (Fox *et al*., 2018; Benini *et al*., 2015), metabolites (Lavin *et al*., 2025), microRNAs (Nair *et al*., 2020), and extracellular vesicles (Rigamonti *et al*., 2019). In turn, many of these individual exercise factors have been identified as having anticancer properties, as demonstrated by their inhibitory effects when directly administered to colorectal cancer cell lines (Ozerklig *et al*., 2025; Orange *et al*., 2022; Wang *et al*., 2022). Given these distinct responses, it is plausible that sex and CRF differences may exist in the effects of exercise on cancer cell outcomes, although this remains to be established. Identifying individual characteristics that play a role in the variability of the effects of exercise on cancer cells holds important implications for supporting a movement from generic exercise prescription towards a precision exercise approach to maximise the potential of exercise in colorectal cancer risk reduction.

The current Acute Exercise Response on Behaviour in Colorectal Cancer Cells (AEROBIC) study assessed the effects of human serum conditioned by acute moderate-intensity aerobic exercise on colorectal cancer cell viability in vitro. The primary aim was to investigate whether sex and CRF modulate the presence and/or magnitude of cancer-suppressive effects of acute aerobic exercise serum. The influence of these individual characteristics at rest on colorectal cancer cell viability in vitro was also examined.

## Methods

### Ethical approval and participants

The study was approved by Swansea University College of Engineering Ethics Committee (ref: RK_30-07-21b) and complied with the Declaration of Helsinki (2013), with the exception of clause 35 (preregistration in a publicly accessible database). Forty-one healthy male (*n =* 28) and female (*n* = 13) volunteers gave written informed consent to participate in the study between September 2021 and June 2022. Exclusion criteria included age < 18 or > 40 years; a history of cardiovascular, metabolic, pulmonary, or immune-related disease; smoking; pregnancy; acute illness or infection in the 7-days preceding experimental trials; blood pressure ≥ 140/90 mmHg and/or resting heart rate > 100 b·min^−1^ measured at the screening visit; and/or any contraindications to exercise according to the Physical Activity Readiness Questionnaire (Thomas *et al*., 1992).

### Cardiorespiratory fitness assessment and categorisation

CRF was determined by peak oxygen consumption (V̇O_2_peak) during a maximal exercise test on an electronically braked cycle ergometer (Lode Excalibur Sport, Lode BV, Netherlands). Participants were instructed to avoid exercise, alcohol, over-the-counter medication for 24 h, and food/caffeine for 2 h prior to the test. The test began with a 3-min warm-up at 20 W followed by an increasing intensity of 1 W every 3 seconds (20 W·min^−1^) until volitional exhaustion, or until pedal cadence could not be maintained ≥ 50 r·min^−1^ despite strong verbal encouragement. Pulmonary gas exchange was measured continuously using a metabolic cart (Vyntus, Jaeger, Germany). Heart rate (HR; Polar Electro, Kempele, Finland) and rating of perceived exertion (RPE) according to the 15-point Borg Scale (Borg, 1982) were recorded every minute and at the point of volitional exhaustion.

V̇O_2_peak was taken as the highest 15-breath rolling average for V̇O_2_ calculated from breath-by-breath data. From this, linear regression (power output vs. V̇O_2_) was used to calculate individual power outputs equivalent to 65% V̇O_2_peak for the experimental trial (Hwang *et al*., 2020; De Santi *et al*., 2019; Rundqvist *et al*., 2013). To verify the predicted power output, following 15-min seated rest, participants cycled continuously for 4-min at their individually calculated workload to allow for steady state to be reached, with pedal cadence maintained between 70 and 80 r·min^−1^ to match the experimental trial. Pulmonary gas exchange was measured throughout, with V̇O_2_ averaged over the last min of the 4-min stage. If necessary, adjustments were made to the workload, and the protocol repeated until the correct V̇O_2_ was achieved (65 ± 2%).

Participants were stratified into two groups based on predefined sex-specific V̇O_2_peak cut-off values: (1) high CRF (males > 45 ml·kg^−1^·min^−1^; females > 40 ml·kg^−1^·min^−1^); or (2) low CRF (males ≤ 45 ml·kg^−1^·min^−1^; females ≤ 40 ml·kg^−1^·min^−1^).

### Experimental design

The AEROBIC study used a randomised, controlled, counterbalanced crossover design (Figure 1). Participants completed two experimental trials: (1) 30-min moderate-intensity aerobic exercise (Exercise); and (2) 30-min seated rest (Control), separated by at least five days.

**Figure 1.**
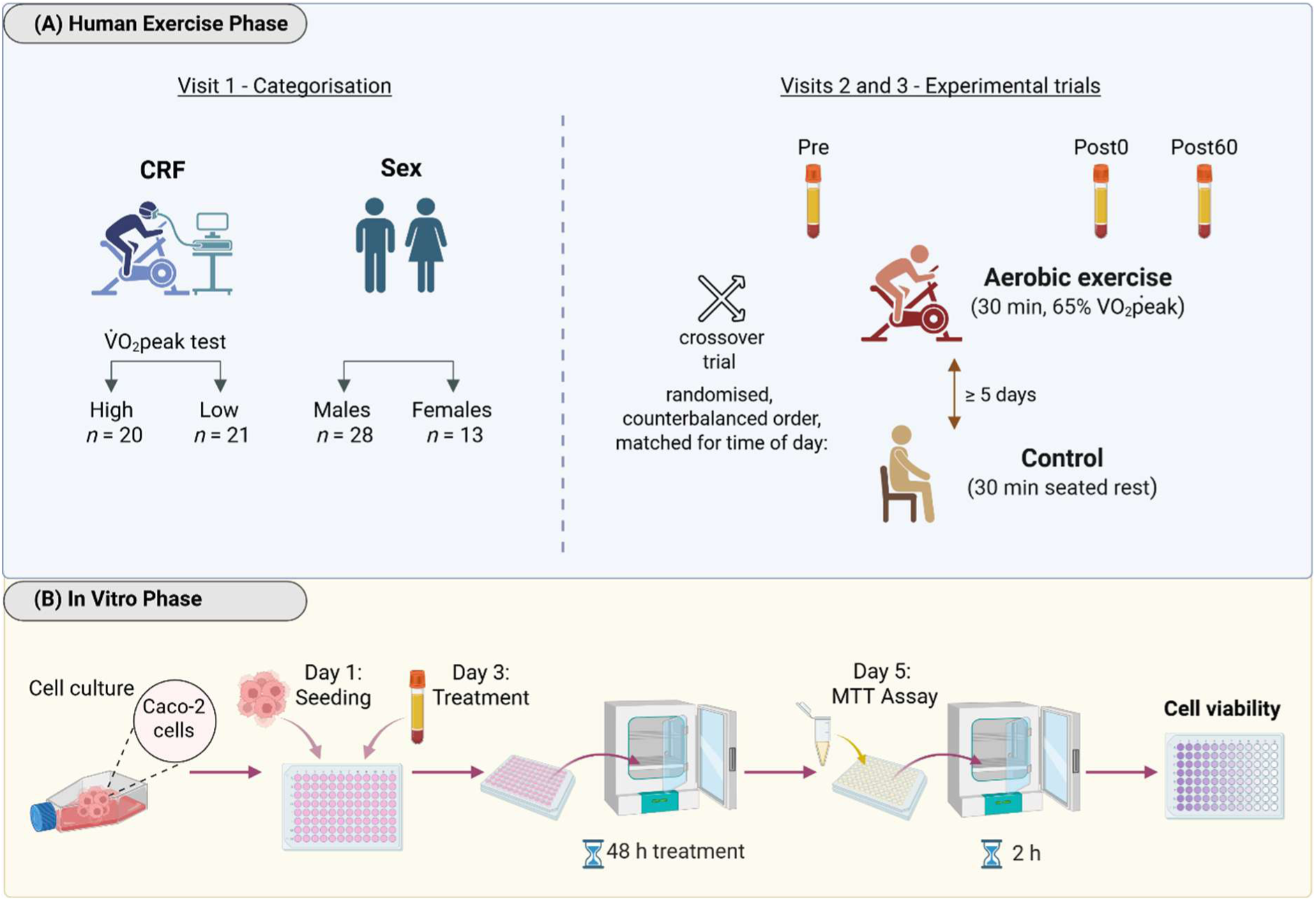
Overview of study design. CRF, cardiorespiratory fitness; V̇O_2_peak, peak oxygen consumption; Post0 and Post60, immediately after and 60-min after exercise or seated rest.

All trials started between 07:00 – 10:00 and were matched for time of day (± 10-min) within participants to control for diurnal variation. In females, trials were matched for menstrual cycle phase (follicular or luteal) within participants, with 62% of trials completed in the follicular phase, as determined by a self-report menstrual cycle questionnaire. Prior to both trials, participants were instructed to refrain from exercise for ≥ 72 h to avoid any residual effects on circulating factors. Participants were also instructed to avoid alcohol and over-the-counter medication for 48 h, and to record and replicate a 24 h food diary prior to each trial. There were no significant differences in total daily energy, carbohydrate, protein, or fat intake within participants for the 24 h preceding trials, analysed using commercially available nutritional analysis software (Dietplan7, Forestfield Software Ltd., UK). On the trial morning, participants arrived after an overnight fast (12 h) and in a euhydrated state having consumed water prescribed proportional to 35 ml·kg^−1^·d^−1^ *pro rata* for the 90-min pre-trial period.

### Acute aerobic exercise and seated rest control trials

On arrival for each trial, stature and body mass were recorded and participants sat quietly at rest for 15 min prior to the collection of the baseline venous blood sample. The aerobic exercise trial consisted of 30-min continuous cycling at moderate-intensity (65% V̇O_2_peak), performed on an electronically braked cycle ergometer (Lode Excalibur Sport, Lode BV, Netherlands) with pedal cadence maintained between 70 and 80 r·min^−1^. Breath-by-breath pulmonary gas exchange was measured continuously (Vyntus, Jaeger, Germany). HR was measured continuously as a secondary objective measure of intensity, and RPE was collected at 5-min intervals using the 6 – 20 Borg Scale (Borg, 1982). For the control trial, participants sat quietly for 30-min under the same environmental conditions with HR measured continuously throughout.

### Serum collection

Blood samples were collected before (pre), immediately after (post0), and 60-min after (post60) exercise or seated rest. At each time point, blood was drawn from different sites within the antecubital fossa into two 6 ml dry vacutainer tubes. Samples were allowed to clot for 50-min at room temperature and then immediately centrifuged (1500 *g* for 10-min at 4 °C). The serum supernatant was aliquoted into sterile 1.5 ml tubes and immediately stored at −80 °C until in vitro experiments. All serum processing procedures were carried out in a class II microbiological safety cabinet using aseptic technique.

### Culture of Caco-2 cells

The human colorectal cancer cell line Caco-2 (DSMZ No: ACC 169) was cultured in minimum essential medium (MEM) supplemented with 10% heat-inactivated fetal bovine serum (FBS; Gibco™), 1% glutamine (Invitrogen™), and 1% penicillin-streptomycin (Gibco™). Cells were maintained at 37 °C in a humidified incubator (5% CO_2_) and routinely passaged when ∼80% confluent, with media replaced every 48 h. Briefly, during passaging, cells were harvested using 0.25% trypsin-EDTA, and cell counts and viability were assessed using the trypan blue exclusion assay (Sigma-Aldrich, UK) and LUNA-II™ automated image-based cell counter (Logos Biosystems) before being split into new flasks for further cultivation or seeded for experiments. Cells were passaged at least three times after thawing from liquid nitrogen and used at passage numbers 5 – 16 for experiments. All cell culture and assay procedures were carried out in a laminar class II tissue culture hood using aseptic technique.

### Cell viability assay

The effect of the basal and exercise-conditioned serum samples on cell viability was assessed using the 3-(4,5-dimethylthiazol-2-yl)-2,5-diphenyl-2H-tetrazolium bromide (MTT) assay (Sigma-Aldrich, UK). This colorimetric assay is based on the reduction of MTT tetrazolium salt to a purple formazan by metabolically active viable cells (Riss *et al*., 2016). Cells were seeded in 96-well plates at a density of 5 × 10^4^ cells·well^−1^. Initially, 100 μl·well^−1^ of cell suspension using normal culture medium was added and plates incubated for 48 h to allow for attachment. Next, the medium was replaced with either 100 μl of medium containing 10% human serum from individual participants instead of 10% FBS (treated cells) or normal FBS-containing medium for the three control wells. Two wells with media only (no cells seeded) were also included on each plate as reference blanks. Samples from each individual were used on the same plate to negate inter-plate variability for within-participant comparisons. Only the inner 60 wells were used for treatment, control, or blanks, with media only added to the perimeter wells to avoid the edge effect (Larsson *et al*., 2020). Serum samples were kept on ice during thawing and treatment, and a new 1.5 ml serum aliquot was used per plate (biological replicate) to avoid any freeze-thaw cycles. After adding the samples, cells were incubated for 48 h (37 °C, 5% CO_2_).

After 48 h treatment, media was removed and 50 μl·well^−1^ of MTT solution (0.5 mg/ml) was added, plates wrapped in aluminium foil, and incubated for 2 h (37 °C, 5% CO_2_). The supernatant was then removed, and formazan crystals were solubilised in 50 μl·well^−1^ dimethyl sulfoxide (DMSO) and mixed via orbital shaking for 10-min. Optical density (OD) absorbance was measured immediately at 570 nm using a FLUOStar Omega microplate reader (BMG LabTech, Ortenberg, Germany). For each serum sample (*n* = 232), three independent experiments were performed using cells from different frozen stocks with different passage numbers (biological replicates). Each biological replicate consisted of three technical replicates (i.e. three replicate wells of identically treated cells), resulting in nine measurements for each serum sample, of which a mean value was calculated for statistical analysis to represent *n* = 1.

### Statistical analyses

Multiple linear regression and independent *t* tests (following normality verification using the Shapiro-Wilk test) were used to determine the influence of individual characteristics at rest on cell viability (cross-sectional design, *n* = 41). Linear mixed-effects models (LMEMs) were used to determine differences in cell viability between exercise and control conditions, and whether individual characteristics moderated the effect of exercise on cell viability (interventional design, *n* = 40). Condition with two levels (exercise and control), serum collection time point with three levels (pre, post0, post60), and their interaction were specified as fixed effects, and participants as a random intercept with random slopes for time and/or condition (Bell *et al*., 2019). Sex, CRF, and baseline values were included as covariates. Models were fitted using restricted maximum likelihood estimation. The assumption of normality of model residuals was verified using Q-Q plots. Statistical significance was set at *P* < 0.05. When a significant difference was detected, postestimation pairwise comparisons with Bonferroni adjustments were used to identify the specific location. Statistical analyses were performed using Stata version 18.5 (StataCorp, College Station, TX, USA).

Effect sizes (Cohen’s *d_rm_*) were calculated to quantify the magnitude of any exercise-induced effects, where a negative value reflects a reduction in cell viability, and, therefore, a favourable small (≥ −0.2), moderate (≥ −0.5), or large (≥ −0.8) effect (Cohen, 1988).

Cell viability results are expressed as the mean ± SEM of three independent experiments, unless otherwise stated. OD single technical replicate outliers, identified using the Grubbs’ test at 0.05 alpha level in GraphPad Prism (v10.0.2, GraphPad Software, USA), were removed prior to any cell viability calculations. Any missing data in blood sample time points is reflected in figures and was handled by LMEMs under the missing at random assumption. Participant and exercise characteristics data are presented as mean ± SD, unless otherwise stated.

## Results

### Participant characteristics

Characteristics of the participants stratified by sex and CRF group are reported in Table 1. Importantly, aside from the characteristic of interest, other potential confounding variables, such as age and body mass index, were similar between subgroups (Table 1).

**Table 1.** Participant characteristics by subgroups (*n* = 41).

|  | Total | Sex |  | CRF |  |
| --- | --- | --- | --- | --- | --- |
|  |  | Males | Females | High | Low |
| $n$ | 41 | 28 | 13 | 20 (6 F) | 21 (7 F) |
| Age (y) | $23.0 \pm 4.2$ | $23.1 \pm 4.4$ | $23.0 \pm 3.9$ | $22.5 \pm 3.5$ | $23.6 \pm 4.8$ |
| $\dot{V}O_{2peak}$ | | | | | |
| ($ml \cdot kg^{-1} \cdot min^{-1}$) | $44.1 \pm 10.0$ | $46.6 \pm 9.8$ | $38.7 \pm 8.8$ | $52.0 \pm 6.2$ | $36.6 \pm 6.7$ |
| ($L \cdot min^{-1}$) | $3.0 \pm 0.9$ | $3.4 \pm 0.8$ | $2.2 \pm 0.5$ | $3.6 \pm 0.8$ | $2.5 \pm 0.7$ |
| Body mass (kg) | $69.0 \pm 14.3$ | $74.3 \pm 13.8$ | $57.6 \pm 7.0$ | $68.4 \pm 11.6$ | $69.5 \pm 16.8$ |
| Height (cm) | $174.8 \pm 9.0$ | $178.6 \pm 7.8$ | $166.7 \pm 5.1$ | $174.8 \pm 9.5$ | $174.9 \pm 8.7$ |
| BMI ($kg \cdot m^{-2}$) | $22.4 \pm 3.3$ | $23.1 \pm 3.3$ | $20.7 \pm 2.8$ | $22.2 \pm 2.1$ | $22.5 \pm 4.3$ |

Continuous variables are mean ± SD. *Abbreviations:* CRF; cardiorespiratory fitness; $n$ , sample size; F, female; $\dot{V}O_{2max}$ , maximal oxygen consumption; BMI, body mass index.

### Impact of sex and cardiorespiratory fitness at rest on Caco-2 cell viability

In basal resting samples, no sex differences in Caco-2 cell viability were observed (*P* = 0.90; Figure 2A). Specifically, compared to 10% FBS control, media supplemented with 10% human male and female serum resulted in 95.03 ± 3.27% and 95.69 ± 3.51% viable cells, respectively. Similarly, no baseline differences in cell viability were observed between resting serum from individuals with high (94.54 ± 3.86%) and low CRF (95.92 ± 3.20%; *P* = 0.78; Figure 2B). When treated as a continuous predictor variable, CRF had a negative but non-significant association with Caco-2 cell viability in isolation (β = −0.18, *P* = 0.47, 95% CI −0.69 to 0.32) and in combination with sex (β = −0.20, *P* = 0.47, 95% CI −0.75 to 0.35).

**Figure 2.**
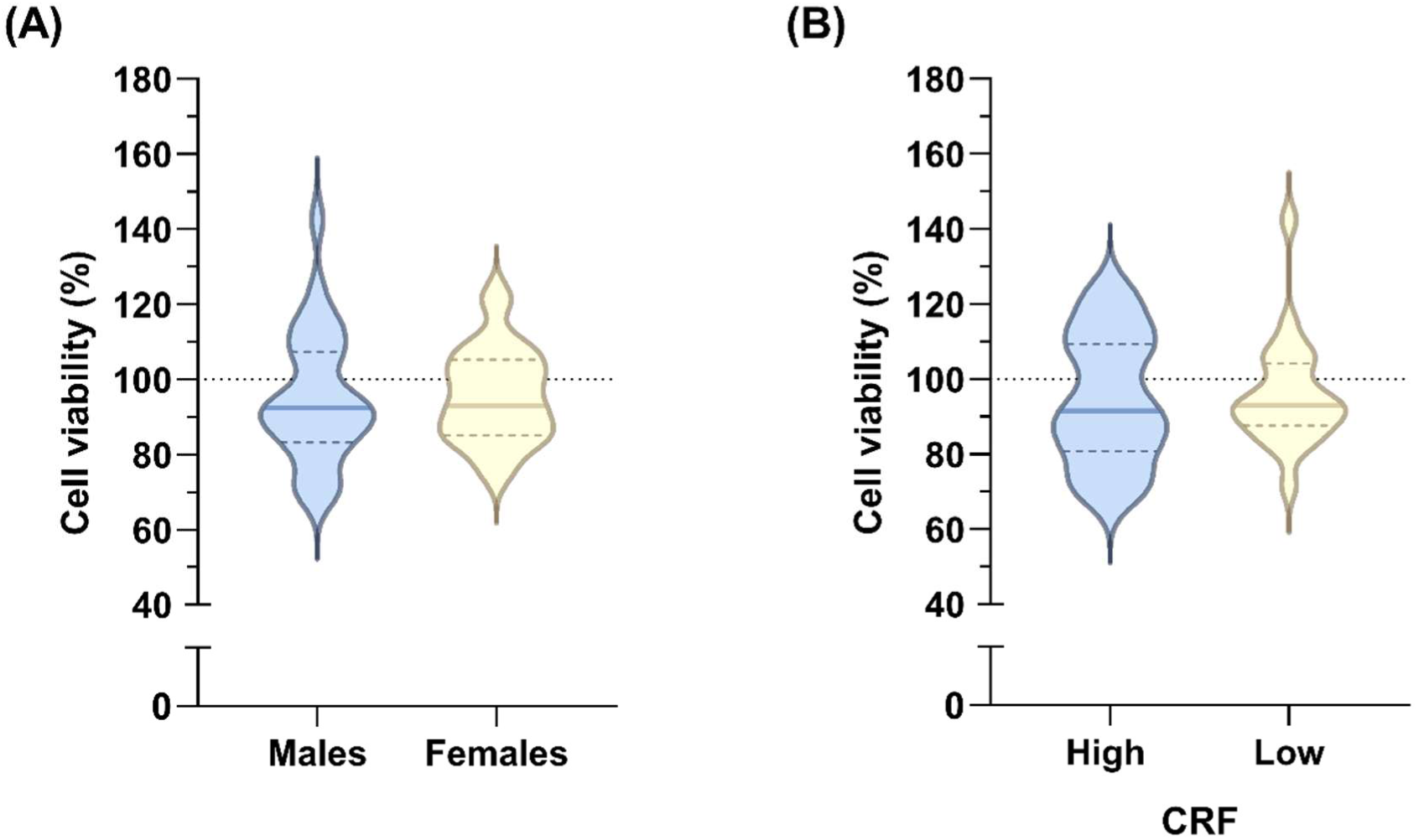
Caco-2 cell viability following treatment with human serum collected at rest from (A) males (*n* = 28) vs. females (*n* = 13); and (B) high (*n* = 20) vs. low (*n* = 21) cardiorespiratory fitness (CRF) groups, compared to FBS control (100% line). Violin plots display the median (central marked line) and 25^th^ and 75^th^ percentiles (dashed lines). Data are representative of three independent experiments, each with three technical replicates, conducted for each individual participants serum (*n* = 41).

### Characteristics of the exercise bout

Forty participants (27 male, 13 female) completed the 30-min cycling bout. The mean V̇O_2_peak was 66.4 ± 5.6% (median 65.3%, IQR 63.5 – 67.2%), indicating high fidelity to the exercise protocol. As secondary measures of exercise intensity, the mean HR was 149 ± 15 b·min^−1^ (75 ± 7% HRmax), the respiratory exchange ratio was 0.91 ± 0.04, and RPE was 13 ± 2. A positive affective valence during the exercise was reported (2 ± 2).

### Cardiorespiratory fitness modulates the colorectal cancer-suppressive effects of acute aerobic exercise serum

When examining the participant population as a whole (*n* = 40), the viability of Caco-2 cells decreased from 94.82 ± 2.65% with pre-exercise serum to 92.47 ± 2.68% with post0 (−2.35%) and 89.14 ± 2.47% with post60 exercise serum (−5.68%; Figure 3). In contrast, during the control trial, cell viability increased by 2.30 ± 1.25% and 2.61 ± 2.16% at post0 and post60, respectively, relative to pre-control serum (Figure 3). Overall, this resulted in a moderate and significant anti-viability effect of exercise serum at post0 (mean difference = −4.65%, β = −4.56, *P* = 0.010, 95% CI −8.04 to −1.08; Cohen’s *d_rm_* = −0.54) and post60 (mean difference = −8.29%, β = −5.84, *P* = 0.001, 95% CI −9.44 to −2.25; Cohen’s *d_rm_* = −0.70) compared to no-exercise control serum.

**Figure 3.**
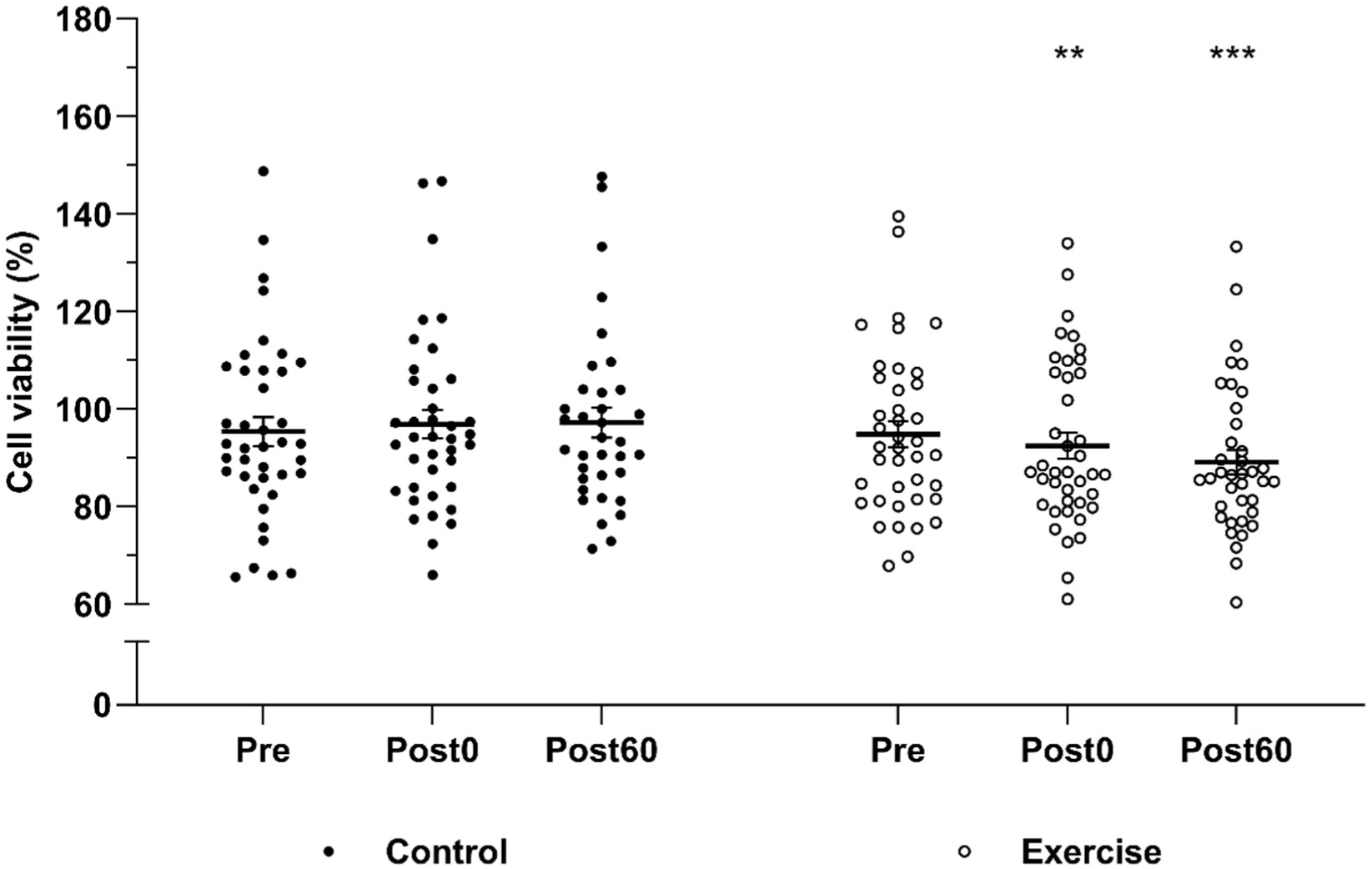
Overall effect of acute moderate-intensity aerobic exercise serum on Caco-2 cell viability with all participants combined (*n* = 40). Comparison of viable cells after 48 h treatment with human serum collected before (pre), immediately after (post0), and 60-min after (post60) exercise and a time-matched no-exercise control trial. Individual data points represent the mean of three independent experiments, each with three technical replicates, for each participant. Bars represent the mean ± SEM. \*\**P* < 0.01 and \*\*\**P* < 0.001, significant condition-time interaction effect (Linear mixed-effects model).

Stratification of participants by V̇O_2_peak revealed that these exercise-induced alterations in Caco-2 cell viability were, at least in part, modulated by CRF (Figure 4). At post0, exercise serum significantly reduced cell viability compared to no-exercise control serum in the high (mean difference = −6.26%, β = −6.51, *P* = 0.027, 95% CI −12.29 to −0.74; Cohen’s *d_rm_* = −0.70) but not low CRF group (mean difference = −3.19%, β = −2.62, *P* = 0.21, 95% CI −6.67 to 1.44; Cohen’s *d_rm_* = −0.39). Similarly, at post60, exercise serum significantly reduced cell viability compared to no-exercise control serum in the high (mean difference −8.86%, β = −7.80, *P* = 0.010, 95% CI −13.71 to −1.89; Cohen’s *d_rm_* = −0.65) but not low CRF group (mean difference −7.71%, β = −3.73, *P* = 0.085, 95% CI −7.98 to 0.52; Cohen’s *d_rm_* = −0.74). Specifically, in the high CRF group, exercise serum reduced cell viability by −4.03 ± 2.19% and −6.33 ± 1.91% at post0 and post60, respectively, compared to pre-exercise serum. In contrast, no-exercise control serum led to a 2.23 ± 1.84% and 2.52 ± 3.53% increase in cell viability at post0 and post60, respectively, compared to pre-control serum (Figure 4B). In the low CRF group, exercise serum reduced cell viability to a lesser extent by −0.83 ± 1.88% and −5.04 ± 1.98% at post0 and post60, respectively, compared to pre-exercise serum, whereas no-exercise control serum resulted in a 2.36 ± 1.76% and 2.67 ± 2.64% increase in cell viability at post0 and post60, respectively, compared to pre-control serum (Figure 4B).

**Figure 4.**
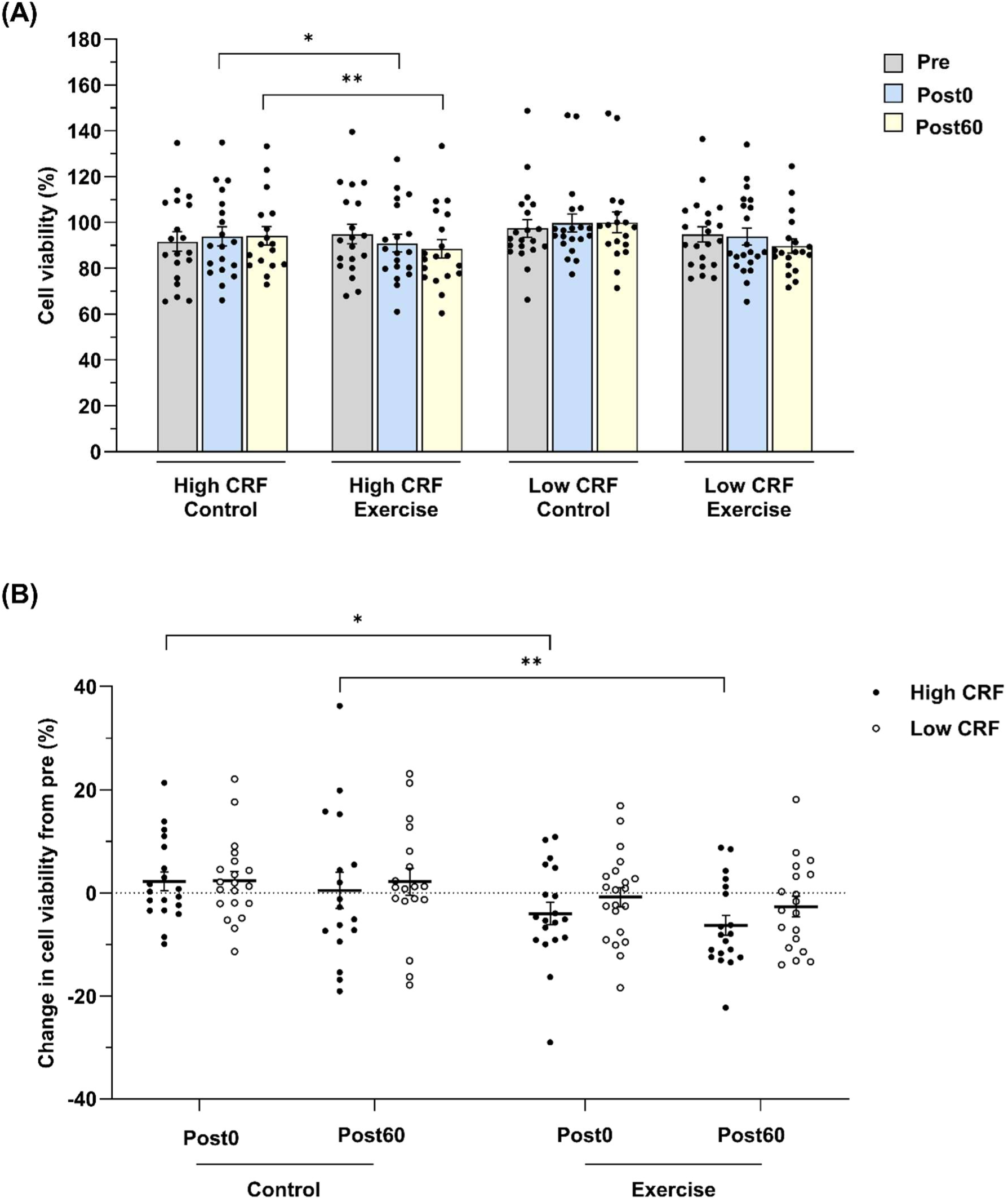
The influence of cardiorespiratory fitness (CRF) on the anti-viability effects of acute aerobic exercise serum on colorectal cancer cells. (A) Caco-2 cell viability after 48 h treatment with human serum collected before (pre), immediately after (post0), and 60-min after (post60) exercise and a time-matched no-exercise control trial from individuals with high (*n* = 19) vs. low (*n* = 21) CRF. (B) Change in cell viability in exercise and control trials. Individual data points represent the mean of three independent experiments, each with three technical replicates, for each participant. Bars represent the mean ± SEM. \**P* < 0.05 and \*\**P* < 0.01, significant condition-time interaction effect (Linear mixed-effects model).

### Sex differences in the effects of acute aerobic exercise serum on Caco-2 cell viability

There was a sex-dependent moderation of the exercise treatment effect on cell viability (Figure 5). Specifically, at post0, male exercise serum significantly reduced cell viability compared to no-exercise control serum (mean difference = −5.59%, β = −5.41, *P* = 0.014, 95% CI −9.71 to −1.11; Cohen’s *d_rm_* = −0.64), whereas female exercise serum had a small and non-statistically significant effect (mean difference = −2.68%, β = 2.57, *P* = 0.39, 95% CI −8.46 to 3.31; Cohen’s *d_rm_* = −0.32). Similarly, at post60, male exercise serum elicited a large and significant reduction in cell viability compared to no-exercise control serum (mean difference = −10.41%, β = −7.64, *P* = 0.001, 95% CI −12.04 to −3.24; Cohen’s *d_rm_* = −0.87), while the effect of female exercise serum remained small and non-statistically significant (mean difference = −3.46%, β = −1.50, *P* = 0.63, 95% CI −7.68 to 4.68; Cohen’s *d_rm_* = −0.30). Within the exercise condition, at post0, cell viability decreased by −3.28 ± 1.76% in males and −0.41 ± 2.50% in females compared to pre-exercise serum (Figure 5B). At post60, male and female exercise serum reduced cell viability by −6.30 ± 1.62% and −4.42 ± 2.77%, respectively, compared to pre-exercise serum (Figure 5B).

**Figure 5.**
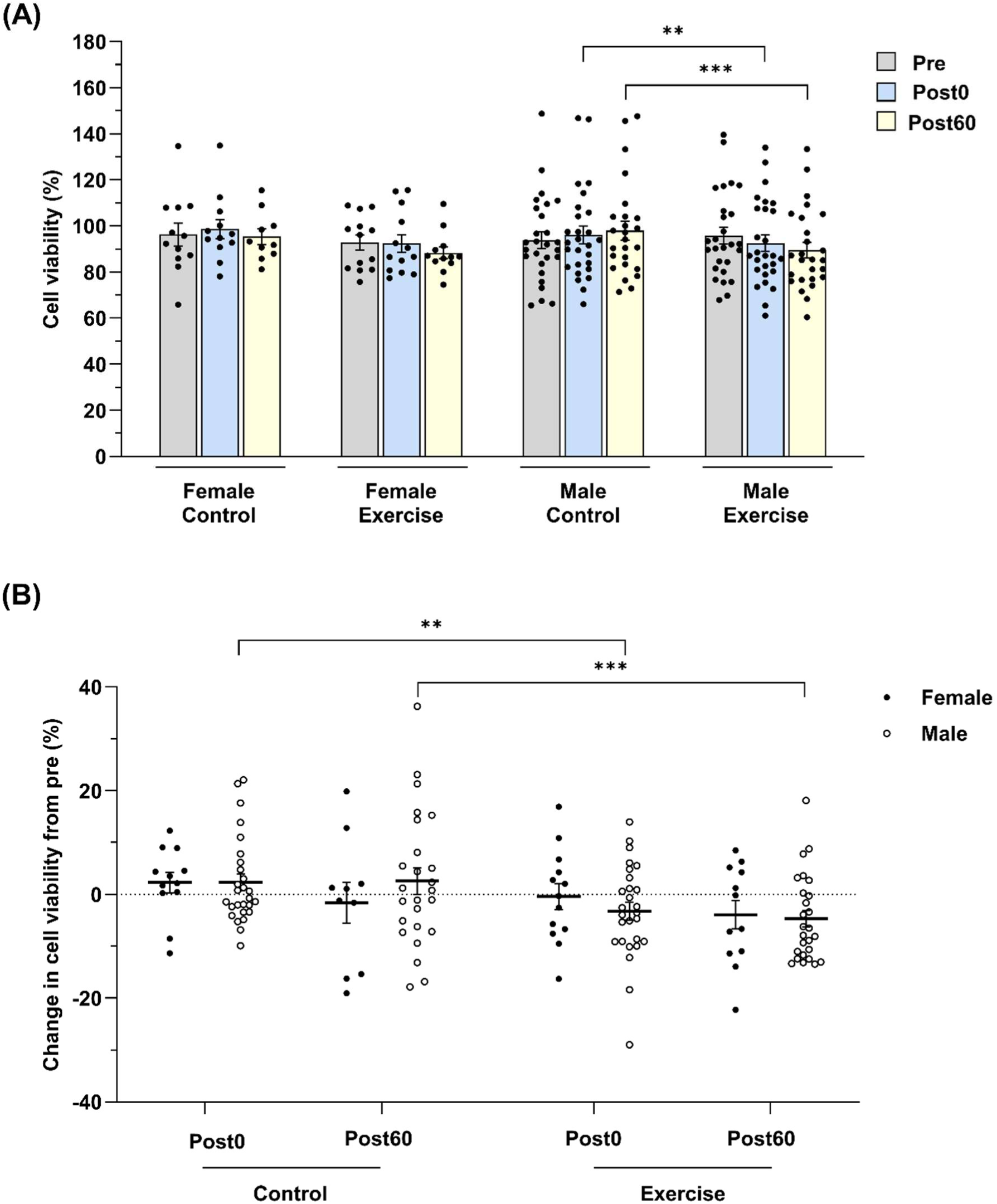
The influence of sex on the anti-viability effects of acute aerobic exercise serum on colorectal cancer cells. (A) Caco-2 cell viability after 48 h treatment with human male (*n* = 27) vs. female (*n* = 13) serum collected before (pre), immediately after (post0), and 60-min after (post60) exercise and a time-matched no-exercise control trial. (B) Change in cell viability in exercise and control trials. Individual data points represent the mean of three independent experiments, each with three technical replicates, for each participant. Bars represent the mean ± SEM. \*\**P* < 0.01 and \*\*\**P* < 0.001, significant condition-time interaction effect (Linear mixed-effects model).

## Discussion

By applying human serum from participant populations with different characteristics as treatment stimuli, this study provides the first insights into the interplay between individual characteristics and acute exercise on colorectal cancer cell viability in vitro. Our results indicate an acute aerobic exercise-induced reduction in Caco-2 cell viability, which was modulated by sex and CRF. Specifically, significant anti-viability effects were present following treatment with exercise serum from individuals with high but not low CRF, and from males but not females. Therefore, it can be proposed that the participant population from which exercise serum is derived may be vital in determining its anticancer potential. This may have implications to help identify populations most likely to benefit from a given exercise dose and increase the precision of exercise recommendations to maximise the potential of exercise in colorectal cancer risk reduction.

The current findings regarding the modulatory role of CRF suggest for the first time that higher CRF levels are associated with distinct changes in the circulating milieu in response to exercise, which promotes a more potent anticancer environment. As serum represents the collective exercise factor milieu, further mechanistic studies are required to distinguish which specific exercise factors mediate the reduction in cell viability. While many exercise factors are typically secreted to a greater extent in individuals unaccustomed to exercise (Ducharme *et al*., 2025; Croft *et al*., 2009), some appear to have a more pronounced response in trained individuals. One example is the myokine irisin, for which the increases immediately post-exercise were reported to be nearly two-fold greater in fit compared to unfit individuals in a recent meta-analysis (Fox *et al*., 2018). This is of particular interest as irisin has been suggested to play a protective role against colorectal cancer. For example, one study reported that individuals with high serum irisin levels had a 78% lower colorectal cancer risk than those with low levels (Zhu *et al*., 2018). In vitro studies have also indicated that direct administration of recombinant irisin may exhibit anticancer properties in Caco-2 cells by regulating inflammatory and apoptotic pathways (Ozturk *et al*., 2026).

In resting samples, our findings align with a previous study which found no differences in the viability of the prostate cancer cell line LNCaP when treated with plasma derived from healthy young males with high (V̇O_2_peak ≥ 45 ml·kg^−1^·min^−1^) compared to low fitness (Peres *et al*., 2022). The larger sample size in the present study extend these findings, which were limited by small group sizes (*n* = 5; Peres *et al*., 2022). From this, it can be suggested that any differences in the basal circulating profile associated with higher compared to lower CRF, at least in otherwise healthy individuals, does not appear to exert direct beneficial bioactive effects in cancer cells in vitro. Conversely, others have reported that resting serum/plasma obtained from healthy young males and chronic lymphocytic leukaemia patients with higher V̇O_2_peak resulted in lower LNCaP and OSU-CLL cell viability, respectively, compared to those with lower V̇O_2_peak (Sitlinger *et al*., 2021; Sridhar *et al*., 2017). Therefore, given the limited number of studies and mixed results within and between cancer types, the influence of CRF on basal serum-stimulated cancer cell viability requires further investigation.

At rest, there were no sex differences in the effect of human serum on Caco-2 cell viability. These results may be considered contrary to expectations, given the sexual dimorphism reported in colorectal cancer (Rodríguez-Santiago *et al*., 2024). Previous in vitro studies have demonstrated that direct administration of female sex hormones, such as 17β-estradiol (E2), reduces the viability of multiple colorectal cancer cell lines (Hsu *et al*., 2014). In contrast, androgens are associated with more varied results and may both promote or inhibit cell viability depending on the concentration (Roshan *et al*., 2016). However, when examined in isolation, these simplified stimuli are devoid of interactions with other bioactive molecules and are sometimes administered at supraphysiological concentrations. As a result, translatability may be limited in comparison to the application of human serum, which better represents the totality of the circulating environment in vivo (Cela *et al*., 2025).

In contrast to basal serum, we found that exercise-conditioned serum exhibited sex-specific effects. Although the temporal pattern in the effect was similar, with both male and female exercise serum eliciting a greater mean reduction in cell viability at post60 than at post0, the magnitude of the effect was more pronounced, and only statistically significant, in males at both time points. These findings align with previous studies reporting sex differences in the acute systemic response to exercise (Lavin *et al*., 2025; Vislocky *et al*., 2008), and importantly, indicate that these translate to varying therapeutic effectiveness against Caco-2 cells. As one possible example, the most prominent exercise-responsive myokine, interleukin-6 (IL-6), has been suggested to transiently increase at transcriptional and circulating levels to a greater extent in males than in females (Abbasi *et al*., 2016; Benini *et al*., 2015). In turn, IL-6 has been identified as a potential factor mediating the anti-viability effects of acute exercise serum on colorectal cancer cells through a reduction in DNA damage (Orange *et al*., 2022). As the current study is the first to directly compare male- and female-derived exercise serum across any cancer type, these sex-specific findings indicate that the prior evidence of a beneficial inhibitory effect of acute exercise serum on colorectal (Orange *et al*., 2022; Devin *et al*., 2019) and lung (Kurgan *et al*., 2017) cancer cell lines in studies that included males only cannot be extrapolated to females. This underscores the need for future research examining the effects of exercise-conditioned serum on colorectal cancer cells, as well as other non-sex-specific cancers, to prioritise the inclusion of both sexes.

A limitation of our study is that only one human cancer cell line was tested, which was male-derived. Matching the sex origin of the cell line with the human serum being tested may be important due to differences in sex chromosome gene expression, potentially leading to different cellular responses to external stimuli (Park *et al*., 2024). For instance, treatment with chemotherapeutic agents (e.g. 5-fluorouracil) has been shown to inhibit cell viability to a greater extent in female-derived than in male-derived colorectal cancer cell lines (Lim *et al*., 2019). Therefore, it would be advantageous for future studies to assess the effects of female exercise serum on colorectal cancer cell lines derived from female patients, such as HT-29. Additionally, our study population included young (18 – 40 y) healthy adults. Although the incidence rates of early-onset colorectal cancer are rising in individuals aged < 50 years (Berrington de Gonzalez *et al*., 2025), older age remains the primary risk factor. As age may influence the acute exerkine response to exercise (MacNeil *et al*., 2021), the representativeness of the findings to populations most at risk of colorectal cancer should be considered a limitation.

## Conclusion

In conclusion, acute aerobic exercise serum can have suppressive effects on colorectal cancer cell viability in vitro. However, these effects appear to be modulated by CRF and sex, with reduced Caco-2 cell viability evident when treated with exercise serum from individuals with high but not low CRF, and from males but not females. These findings suggest that the characteristics of the individuals from which the exercise serum is derived play an important role in determining its anticancer properties, with potential implications for increasing the precision of exercise recommendations for colorectal cancer risk reduction. Future research should aim to characterise the distinct exercise-induced changes in the circulating milieu between participant populations to identify the specific exercise factors mediating the effect.

## Author contributions

**Rachael Kemp:** conceptualisation, methodology, investigation (human and in vitro), data curation, formal analysis, visualisation, project administration, resources, writing – original draft, writing – review and editing. **Ume-Kulsoom Shah:** methodology (in vitro), resources. **Sarah Alwan:** investigation (human). **David Oyler:** investigation (human). **Gareth Jenkins:** methodology (in vitro), resources. **Kelly Mackintosh:** supervision, writing – review and editing. **Melitta McNarry:** conceptualisation, supervision, writing – review and editing. All authors approved the final version of the manuscript and agree to be accountable for all aspects of the work. All persons designated as authors qualify for authorship, and all those who qualify for authorship are listed.

## Competing interests

The authors declare no competing interests.

## Data availability

All data generated during this study are included in the article. The raw unprocessed version of data are available from the corresponding author upon request.

## Funding

This study received no grant from any funding agency in the public, commercial, or not-for-profit sectors.

## Notes

### Competing Interest Statement

The authors have declared no competing interest.

